# Variational Autoencoders for Generating Synthetic Tractography-Based Bundle Templates in a Low-Data Setting

**DOI:** 10.1101/2023.02.24.529954

**Authors:** Yixue Feng, Bramsh Q. Chandio, Sophia I. Thomopoulos, Tamoghna Chattopadhyay, Paul M. Thompson

## Abstract

White matter tracts generated from whole brain tractography are often processed using automatic segmentation methods with standard atlases. Atlases are generated from hundreds of subjects, which becomes time-consuming to create and difficult to apply to all populations. In this study, we extended our prior work on using a deep generative model a Convolutional Variational Autoencoder - to map complex and data-intensive streamlines to a low-dimensional latent space given a limited sample size of 50 subjects from the ADNI3 dataset, to generate synthetic population-specific bundle templates using Kernel Density Estimation (KDE) on streamline embeddings. We conducted a quantitative shape analysis by calculating bundle shape metrics, and found that our bundle templates better capture the shape distribution of the bundles than the atlas data used in the original segmentation derived from young healthy adults. We further demonstrated the use of our framework for direct bundle segmentation from whole-brain tractograms.

## I. INTRODUCTION

Diffusion MRI[1], [2] and fiber tractography allow for in-vivo reconstruction of white matter tracts [3], [4] and the study of microstructural fiber integrity [5] and structural connectivity in the brain. Tractography can also be used to study psychiatric and neurodegenerative diseases such as Alzheimer’s disease [6], [7]. Given the large number of streamlines generated in whole brain tractograms and numerous fiber bundles of interest, various bundle segmentation methods [8], [9], [10] have been developed to enable automated fiber bundle analyses that internally rely on standard tractography atlas(es) [11], [12]. Atlases are often computed from hundreds of subjects and enable alignment across subjects; they can also be used as templates to assist in bundle segmentation and along-tract analysis [7], [13]. However, even for the same bundles of interest, there is no gold standard for how to define them, and there are often large variations in the resulting segmentations across methods [14]. In addition, segmented bundles can still contain a high proportion of false positive streamlines [15], [16], making it difficult to directly conduct group comparisons.

Standard atlases are often derived from young, healthy populations and they may not be useful for all populations. Spencer *et al*. [17] created age-specific white matter atlases for young children aged 6-8 years old, motivated by the fact that there are developmental changes and adult brain atlases may not be representative of younger populations. Publicly available white matter atlases have also been created for neonates and 1-2 year old populations by Short *et al*. [18]. In addition, atlases sometimes require tens to hundreds of subjects, and those manually segmented by trained neuroanatomists results in lower sample size, overall making the process time consuming.

Deep generative models have gained popularity in recent years, in computer vision for text-to-image generation [19], [20] and in medical imaging to create realistic synthetic images [21]. They can learn a compact representation for complex data distributions using deep neural networks and can be used to generate new realistic samples in both brain imaging and musculo-skeletal radiology [22]. Prior studies have investigated the use of autoencoders, an architecture that encodes high dimensional data into a low dimensional latent space and optimizes the mapping to minimize data reconstruction error, on tractography data. FINTA [23], GESTA [24] ad FIESTA [25] uses streamline-based autoencoders to learn from tractogram(s) for streamline filtering, generative sampling in the latent space and tractogram segmentation. However, training on whole-brain tractograms which contains a large amount of streamlines makes it more difficult to capture variations across more training subjects. In the generative sampling tasks, the implausible streamlines in tractograms used for training can effect what the autoencoder learns as valid streamline locations. Lizarraga *et al*. proposed StreamNet [26], which uses Wasserstein autoencoders (WAE) to learn from bundles directly instead of individual streamlines and generate model bundles. Using subsampled streamlines from bundles comes at the cost of losing shape information of the entire bundle, and mapping one bundle to one point in the embedding space is limited for segmentation and filtering tasks.

In our previous work [27], we used a convolutional variational autoencoder (ConvVAE) to learn low-dimensional embeddings from high-dimensional streamlines, and we used the flexibility of VAE to compute along-tract structural anomalies for group comparisons. We found that ConvVAE was able to preserve inter-streamline and inter-bundle distances well using an embedding space of only 6 dimensions. In addition, the bottleneck in the encoder-decoder was able to denoise and “smooth” streamlines in reconstruction while retaining their location, shape, and orientation. In this study, we further investigate the use of a VAE to capture the distribution of bundle shapes in its latent space with Kernel Density Estimation (KDE). We show that our method can generate synthetic bundle templates from a limited sample size of 50 subjects from the Alzheimer’s Disease Neuroimaging Initiative (ADNI) data, and better captures the population-specific bundle shapes compared to the atlas data that was created from young healthy controls from the Human Connectome Project [28]. We also demonstrate that the estimated kernel density can perform bundle segmentation directly from the whole brain tractograms.

## II. METHOD

### A. Data Processing

We analyzed 3D diffusion MRI data of the brain from 141 subjects - 87 cognitively normal controls (CN), 44 with mild cognitive impairment (MCI), and 10 with dementia (AD) - from the Alzheimer’s Disease Neuroimaging Initiative (ADNI) [29] (age: 55-91 years, 80F, 61M). 3D multi-shell diffusion MRI was acquired from the participants on 3T Siemens scanners, where each scan contains 127 volumes per subject; 13 non-diffusion-weighted *b*_0_ volumes, 6 *b*=500, 48 *b*=1,000 and 60 *b*=2,000 s/mm^2^ volumes with a voxel size of 2.0×2.0×2.0 mm. dMRI volumes were preprocessed using the ADNI3 dMRI protocol, correcting for artifacts including noise [30], [31], [32], [33], Gibbs ringing [34], eddy currents, bias field inhomogeneity, and echo-planar imaging distortions [35], [36], [37]. Whole-brain tractography was calculated using multi-shell multi-tissue constrained spherical deconvolution (MSMT-CSD) [38] and a probabilistic particle filtering tracking algorithm [39]. Auto-calibrated RecoBundles [8], [7] from DiPY [30] was used to extract 30 white matter tracts from all subjects in the MNI space.

### B. Model Design & Training

We trained a variational autoencoder (VAE) on 30 extracted bundles from 50 control subjects, with age ranging from 62 to 86 years (M=71.17, SD=5.81) from the ADNI S127 dataset, with a total of 1,642,183 streamlines. Bundle labels were not used in training, and only the streamlines’ coordinates were passed in as model input, where one streamline is one sample. Each streamline was resampled to 256 equidistant points to allow the use of convolutional layers [23]. All streamlines from each subject were normalized to fit into a standard sphere, where the centroid and radius are calculated from the atlas data (https://figshare.com/articles/dataset/Atlas_of_30_Human_Brain_Bundles_in_MNI_space/12089652)[12] to account for any misalignment between subjects.

The encoder and decoder each had 3 layers of convolution or deconvolution with leaky ReLU activation. Notably, large kernel sizes of 127, 63, and 31 were used in the convolution to better capture long range dependency within streamlines [40]. Empirically, we found that large kernels generate smoother streamlines in reconstruction, compared to smaller kernel sizes (e.g., 3, 5) often used in traditional convolutional neural networks. From our prior work, we found that setting the dimension of the bottleneck layer (*z*) to 6 was optimal for preserving streamline distances in the low-dimensional latent space, and the same dimension was used in the current model setting. The model was trained for 50 epochs with a batch size of 512, using the Adam optimizer with a learning rate of 5e-4 and weight decay of 5e-3. We applied gradient clipping with a max norm of 2. Cyclic annealing [41] was applied to the Kullback–Leibler term to prevent posterior collapse, a common issue in VAE training.

### C Bundle Template Generation

After training the VAE model, we performed inference on all 141 subjects to extract their streamline embeddings *Z*. For each bundle, we trained a kernel density estimator (KDE) on *Z* from 20,000 randomly subsampled streamlines embeddings in the bundle from the 50 subjects used to train the VAE for computational efficiency. We tested 15 bandwidths, ranging from 0.01 to 10, and selected the best KDE based on log-likelihood for downstream analysis.

After training the KDE, we sampled *N* points (ranging from 5,000-10,000) from the fitted KDE to create the bundle template. Despite training and tuning the VAE and KDE, the sampled bundle can still contain noisy samples. We computed the log-likelihood for each streamline in the sample bundles and set the filtering threshold *T* to the mean log-likelihood from the 20,000 streamlines used to train KDE. Depending on the bundle, we can also relax this threshold (e.g., to the 25% percentile of the log-likelihood from the training samples). Selection of *N* and *T* are empirical and depend on the model fit and size of the desired bundle. The filtered generated samples in the low-dimensional latent space were then passed to the VAE decoder to extract the final bundle template. A complete diagram of our experimental design is shown in Figure 1.

**Fig. 1.**
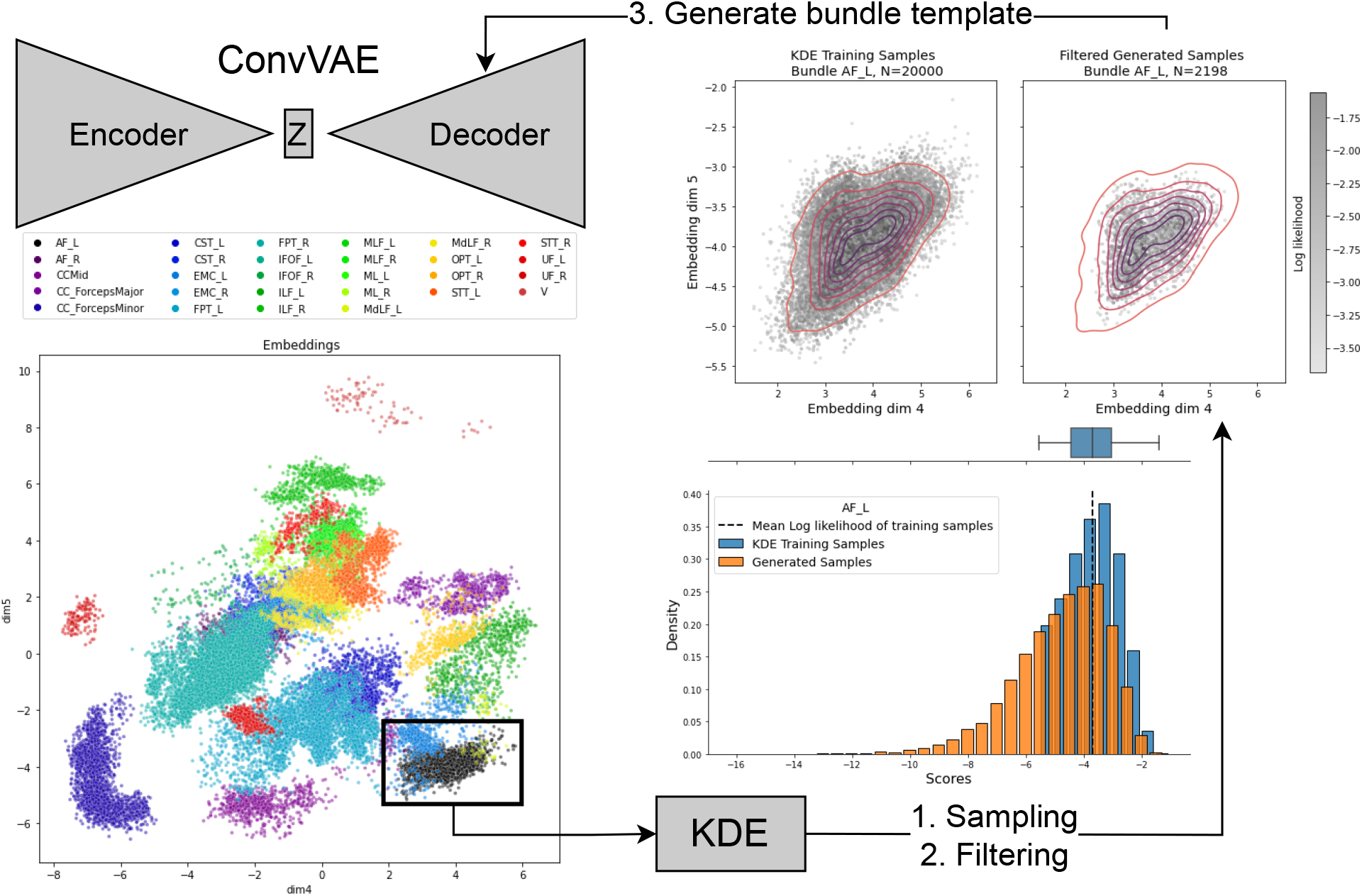
Experiment Diagram. We show example embeddings of an example control (CN) subject not used in training for two of the 6 latent dimensions colored by bundle labels. Note that KDE was fitted on subsampled bundle embeddings from 50 subjects used for ConvVAE training instead of one subject. Sampled points from the fitted KDE are then filtered using a selected threshold calculated from the training sample’s log-likelihood, shown on the lower right plot. The upper right plots show the density plot of the embeddings in 2 dimensions colored by log-likelihood for samples used to fit KDE with contours. The samples generated from KDE after filtering are also shown with the same contour overlayed for comparison. Plots are shown for one bundle, the arcuate fasciculus (AF_L) for demonstration purposes.

### D. Bundle Shape Analysis

To compare our population-specific bundle templates with the atlas data used in bundle segmentation, we first computed the bundle shape similarity (SM) scores [7], [30] between the subject bundles for all subjects excluding those used in training the VAE and KDE, and the template and atlas bundles. We then calculated 6 additional bundle shape metrics - length, span, volume, diameter, surface area, and irregularity, defined in Yeh (2020) [42] to further investigate the features preserved in the low-dimensional embeddings and their distributions learned by the KDE. We expect the bundle templates’ shape to align more closely with the subject bundles than the atlas data. In a downstream task, we demonstrate the use of the fitted KDE to extract bundles directly from whole-brain tractograms.

## III. RESULTS

### A. Bundle Template Evaluation

For a detailed examination of the bundle templates, we show examples for one corpus callosum (corpus callosum forceps minor, CC_ForcepsMinor), one association bundle (left arcuate fasciculus, AF_L), and one projection bundle (left frontopontine tract, FPT_L) in Figure 2. All three template bundles were initially sampled with *N* = 8, 000 from the fitted KDE and decoded by the ConvVAE. The AF_L and CC_ForcepsMinor template were filtered using the mean log-likelihood of the samples used to fit KDE, whereas the FPT_L template was filtered using 25% of the log-likelihood. Without filtering, the template bundle has a larger span and thickness and contains some noisy samples. For FPT_L, selecting a lower threshold yielded results that were more consistent with the atlas and training data considering its size and high number of streamlines per bundle.

**Fig. 2.**
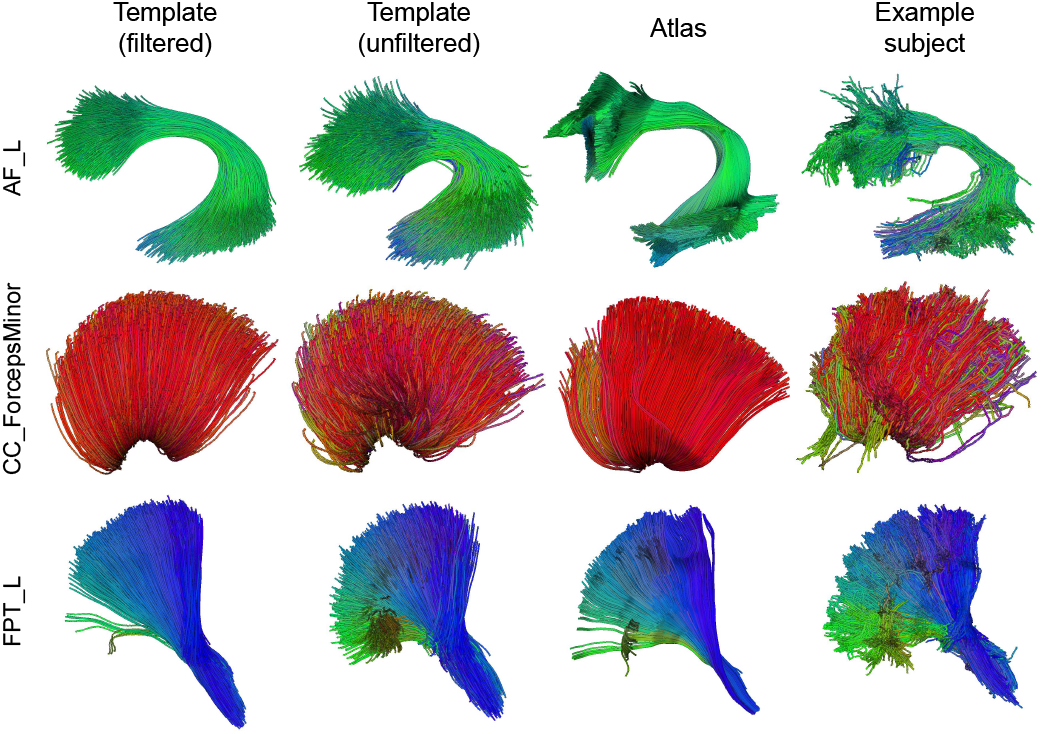
Bundle templates for 3 selected bundles - AF_L, CC_ForcepsMinor, and FPT_L. The filtered, unfiltered bundle templates generated from KDE as described in Section II-C, atlas bundles, and example bundles from a randomly selected control subject are shown from left to right.

In the shape analysis, we computed shape similarity (SM, ranging from 0 to 1, where a higher score indicates greater similarity) of the template and atlas bundles with the subject bundles. SM scores were computed for a total of 91 subjects (CN: 37, MCI: 44, AD: 10), excluding the 50 CN subjects used in training, and we compare these for the 3 diagnostic groups in Figure 3. Overall, SM scores are consistently higher (above 0.8 for most subjects) for the template than the atlas bundles, indicating that the VAE and KDE model is able to capture population-specific bundle shapes despite using the atlas data for bundle segmentation. We also see lower variances in SM for the AF_L and CC_ForcepsMinor templates than the atlas bundles for all 3 groups. Considering that the VAE and KDE are trained on CN subjects, and potential shape anomalies that occur in bundles for subjects with dementia, we see lower shape similarity in the AD group for AF_L and CC_ForcepsMinor, more notable in the atlas data.

**Fig. 3.**
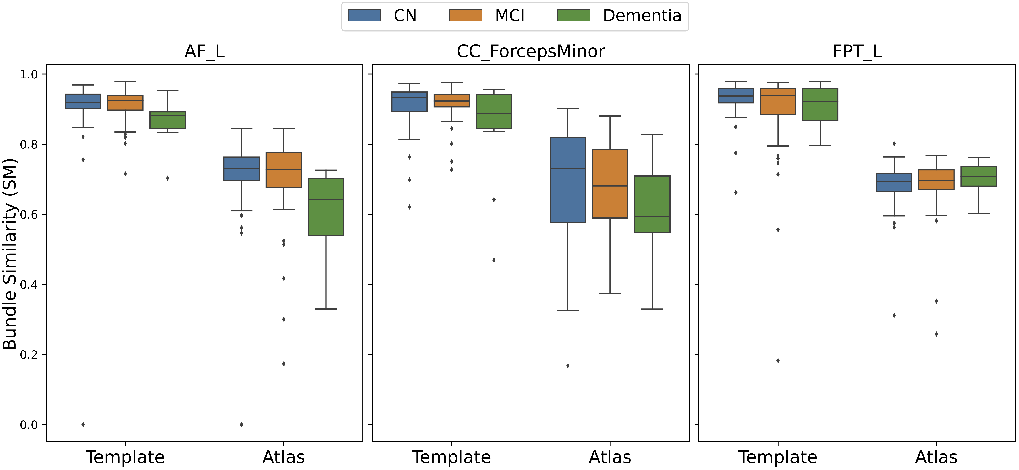
Shape similarity (SM) scores for subject vs. template bundles and subject vs. atlas bundles are shown for 91 subject from the test set across 3 diagnostic groups.

Six shape metrics computed for the template and atlas bundles were also compared with the corresponding subject bundles, see Figure 4, 5, and 6. For all three bundles, the synthetic templates show shorter bundle lengths, whereas the atlas bundles are longer than the subject bundles. The templates align with the subject bundle shape distribution for other metrics, particularly for the span, volume, and surface area. Given the limited sample size used in VAE training and the 20,000 streamlines in estimating kernel density, the templates show comparable results in shape metrics compared to the atlas data.

**Fig. 4.**
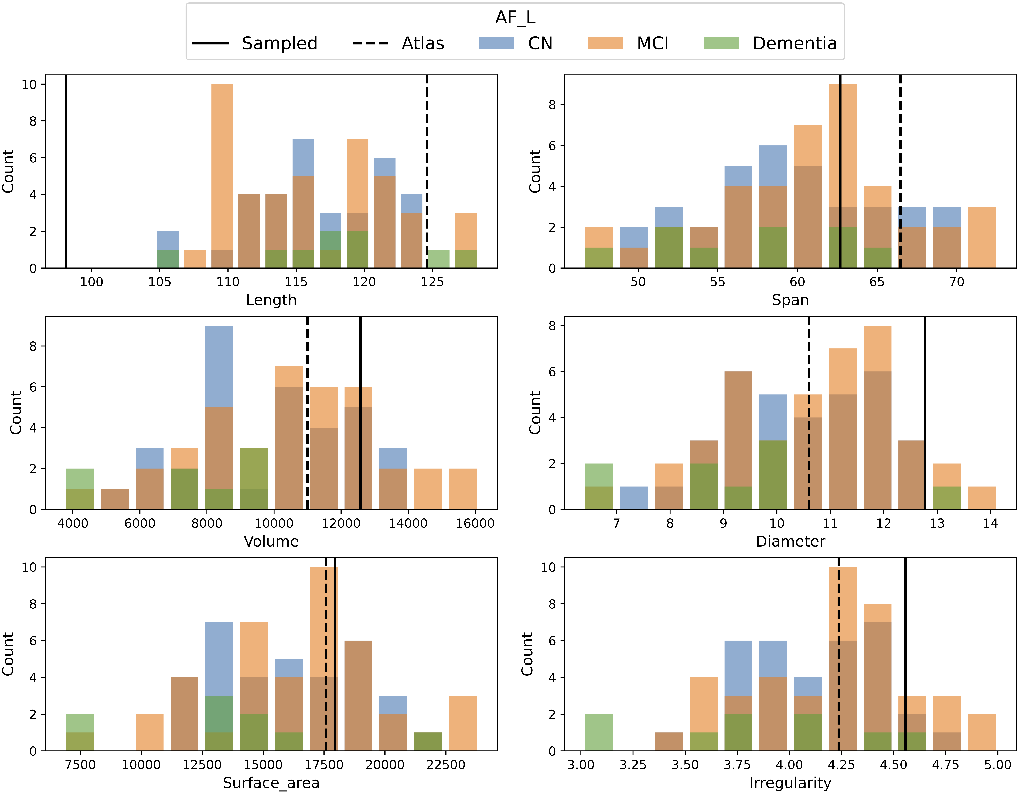
Bundle shape comparison for AF_L.

**Fig. 5.**
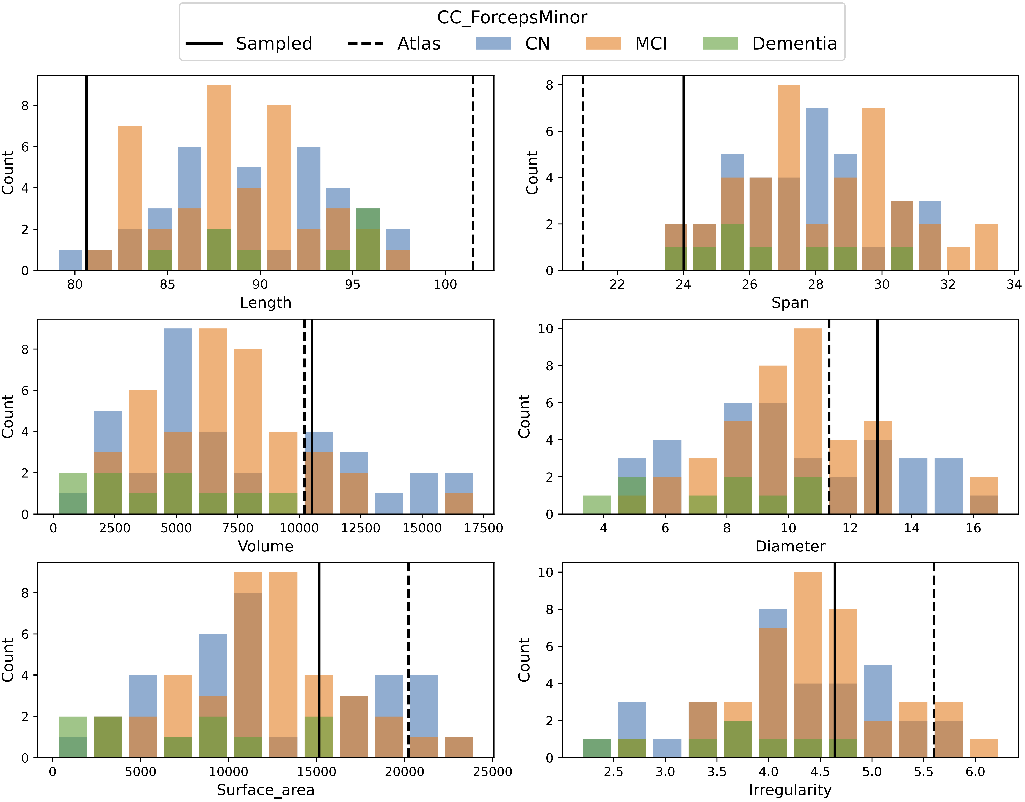
Bundle shape comparison for CC_ForcepsMinor.

**Fig. 6.**
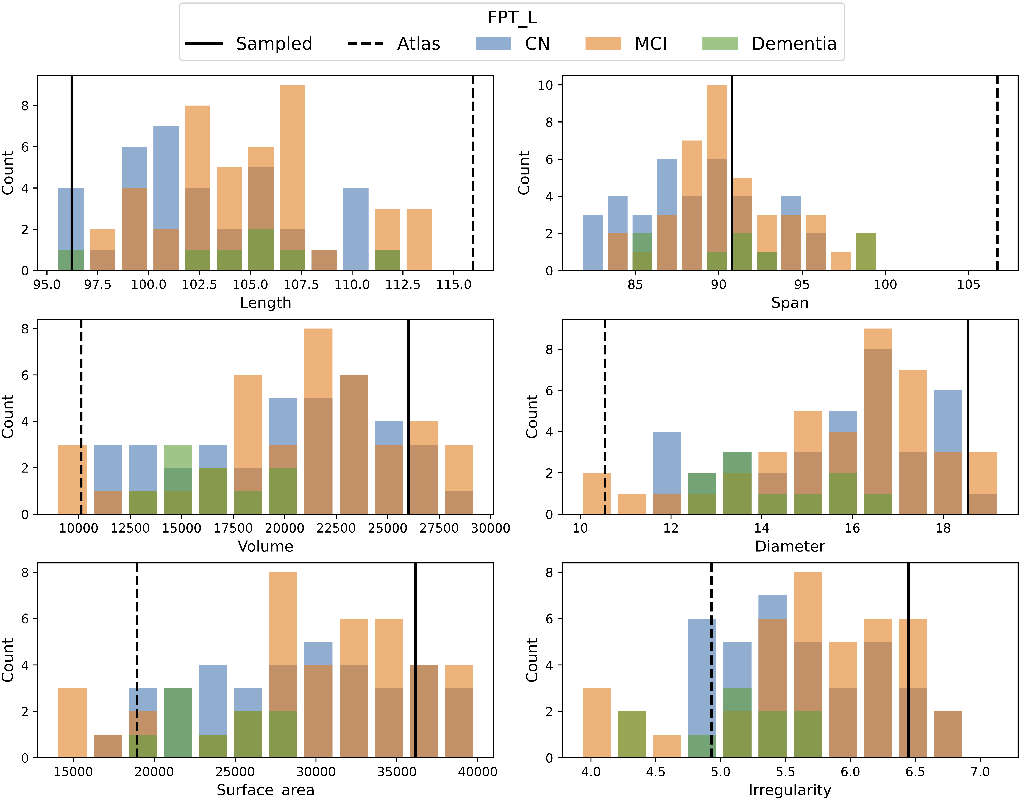
Bundle shape comparison for FPT_L.

### B. Whole-Brain Tractogram Segmentation

In a downstream analysis, we used the fitted KDE for each bundle to perform segmentations directly from whole-brain tractograms. We first perform inference on all streamlines in a given tractogram to extract their embeddings, then calculate the log-likelihood using KDE for each sample. We then filter the results using a more lenient threshold (*Q*_1_ −1.5 × (*Q*_3_ −*Q*_1_) = *Q*_1_ −1.5× *IQR*, calculated from the training samples’ log-likelihood, often used in outlier removal) compared to the previous threshold used in template generation. This threshold can also be adjusted depending on the desired shape of the final bundle. Similarly, the filtered embeddings are then decoded using the ConvVAE decoder. We show bundles segmented using RecoBundles using the atlas as reference bundles and our approach in Figure 7 for one CN and one AD subject not used in training.

**Fig. 7.**
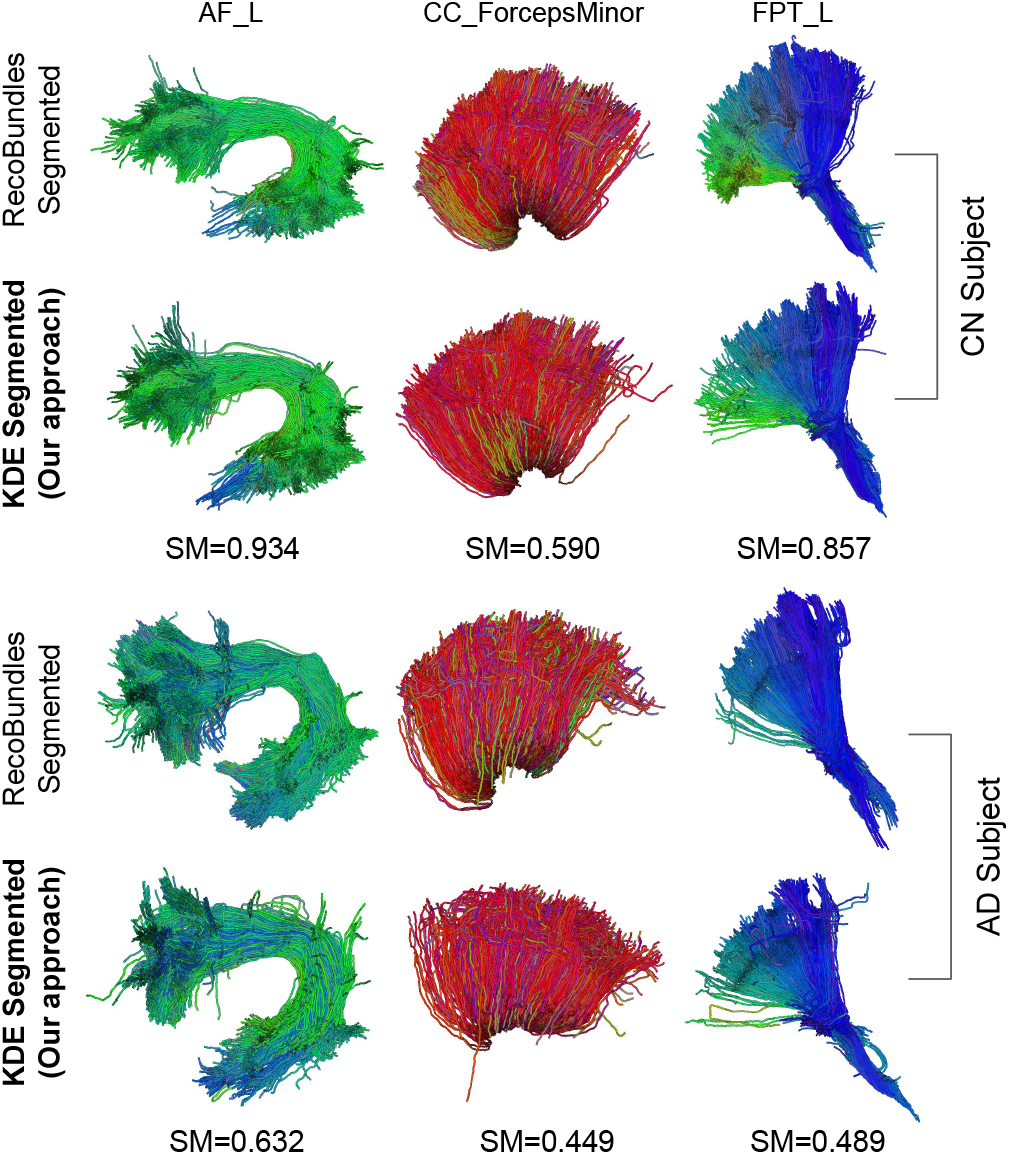
Whole-brain tractogram segmentation results using RecoBundles and KDE for one CN and one AD subject from the test set. Shape similarity (SM) scores between bundles generated using the two approaches are shown below each pair of plots.

The shape similarity (SM) scores between KDE segmented bundles and RecoBundles results are higher for AF_L and FPT_L, and higher in the CN subject for all bundles. Low SM score for CC_ForcepsMinor could be due to the high inter-subject variation and false positive streamlines in the bundles used for VAE training. Overall, the results from our approach largely preserved the shape and location of the bundle, with some slight shape differences. Given that the ConvVAE model inference evaluates each streamline independently, our approach can also be used on incomplete tractograms.

## IV. DISCUSSION

In this study, we proposed to use Kernel Density Estimators (KDE) along with a ConvVAE to learn the shape distributions of bundles, given a limited sample size and bundles that contain false positive streamlines. We showed that our synthetic bundle templates can capture the shape distribution from the training data and show higher shape similarity with the subject bundles. In terms of computation time, the ConvVAE took 42 minutes to train on an NVIDIA Tesla V100 GPU, 3 minutes to fit the KDE for one bundle on an 8-core CPU, and 3 minutes per subject to perform VAE and KDE inference on whole brain tractograms for segmentation. With a trained VAE and KDE, our model is computationally efficient and simple to extend to a large number of subjects with little adaptation.

As demonstrated by the segmentation task in Section <u>III-B,</u> KDE can be used to segment bundles directly from whole-brain tractograms. If we were to select a different segmentation method or atlas, our model can also be adapted to train on various schemas without segmenting a large number of bundles to train on or filtering out the false positive streamlines. While our approach can be used for bundle segmentation, the idea of learning shape distributions with a VAE and KDE has the potential to be extended to more advanced architectures, such as conditional deep generative models (CDGM) [43] where we can use microstructural features such as fractional anisotropy (FA), subject labels and other metadata to supervise the bundle generation process. This opens up more opportunities for using DGM to understand how various neurodegenerative and psychiatric diseases can affect bundle structures, and even model their changes across developmental stages and the human lifespan.

One limitation of the bundle templates generated from KDE is that the bundle lengths are underestimated. This could be due to how the VAE learns the streamline endpoints. With a low dimensional latent space, the VAE is able to learn shape variations along the tract, but wide variations in the bundle endpoints across subjects might not be preserved. For the purpose of bundle segmentation or streamline filtering, regional connectivity of the bundles generated warrants further investigation.

## V. CONCLUSION

In this study, we extended our prior work training a Convolutional Variational Autoencoder (ConvVAE) to extract low-dimensional embeddings from tractography data, to use Kernel Density Estimators (KDE) to learn bundle shapes from a limited sample size of 50 subjects. We generated a synthetic population-specific bundle template from KDE samples and showed that they can better capture the shape distributions on a test set composed of control, MCI, and AD subjects, compared to the atlas bundles, derived from young healthy controls from HCP used in the original segmentation. We further demonstrated the use of KDE for direct bundle segmentation on whole-brain tractograms. Our framework offers an efficient, robust, and flexible approach to understanding the shape of bundle structures, as well as generating synthetic data and potentially aiding in modeling structural abnormalities and changes in bundles in various diseases.

## ACKNOWLEDGMENTS

This work was supported by the National Institutes of Health under the FiberNET project grant, RF1AG057892.

